# Gene losses may contribute to subterranean adaptations in naked mole-rat and blind mole-rat

**DOI:** 10.1101/2021.05.28.446201

**Authors:** Zhi-Zhong Zheng, Rong Hua, Guo-Qiang Xu, Hui Yang, Peng Shi

**Affiliations:** State Key Laboratory of Genetic Resources and Evolution, Kunming Institute of Zoology, Chinese Academy of Sciences, Kunming 650223, China; Kunming College of Life Science, University of Chinese Academy of Sciences, Kunming 650204, China; College of Pharmaceutical Sciences, Soochow University, Suzhou 215006, China; Joint Laboratory of Animal Models for Human Diseases and Drug Development, Soochow University and Kunming Institute of Zoology, Chinese Academy of Sciences, Kunming 650223, China; Center for Excellence in Animal Evolution and Genetics, Chinese Academy of Sciences, Kunming 650223, China; School of Future Technology, University of Chinese Academy of Sciences, Beijing 101408, China

**Author notes:** These authors contributed equally to this work (Zhi-Zhong Zheng, Rong Hua). Correspondence to: Peng Shi, OR, Hui Yang, State Key Laboratory of Genetic Resources and Evolution, Kunming Institute of Zoology, Chinese Academy of Sciences, 32 Jiaochang Donglu, Kunming 650223, China.

**Keywords:** gene losses, naked mole-rat, blind mole-rat, subterranean adaptations

## Abstract

The naked mole-rats (*Heterocephalus glaber*, NMRs) and the blind mole-rats (*Spalax galili*, BMRs) are representative subterranean rodents that evolved many extraordinary traits, including hypoxia tolerance, longevity and cancer resistance. Although a batch of candidate loci responsible for these intriguing traits have been uncovered by genomic studies, many of them are limited to functional modifications of intact genes and little is known about the contributions of other genetic makeups. Here, to address this issue, we focused on gene losses (unitary pseudogenes) and systematically analyzed gene losses in NMRs and BMRs, as well as their respective terrestrial relatives, guinea pigs and rats, in a genome-wide scale. 167, 139, 341 and 112 pseudogenes were identified in NMRs, BMRs, guinea pigs and rats, respectively. Functional enrichment analysis identified 4 shared and 2 species-specific enriched functional groups (EFGs) in subterranean lineages. The pseudogenes in these EFGs might be associated with either regressive (e.g. visual system) or adaptive (e.g. altered DNA damage response) traits. In addition, several pseudogenes including *TNNI3K* and *PDE5A*, might be associated with their specific cardiac features observed in subterranean linages. Furthermore, we observed 20 convergent gene losses in NMRs and BMRs. Given that the functional investigations of these genes are generally scarce, we provided functional evidence that independent loss of *TRIM17* in NMRs and BMRs might be beneficial for neuronal survival under hypoxia, supporting the positive role of eliminating *TRIM17* function in hypoxia adaptation. We also demonstrated that pseudogenes, together with positively selected genes, reinforced subterranean adaptations cooperatively. Overall, our study provides new insights into the molecular underpinnings of subterranean adaptations and highlights the importance of gene losses in mammalian evolution.

## Introduction

Subterranean niches provide mammalian dwellers with highly stressful living conditions that are characterized mainly by extreme hypoxia/hypercapnia, darkness, and food scarcity. Over the course of their evolution for millions of years, subterranean mammals have evolved exquisite physiological and morphological modifications to cope with these challenges (Nevo 1979). Excellent examples of subterranean adaptation are found in naked mole-rats (*Heterocephalus glaber*, NMRs) (Perez, et al. 2009; Seluanov, et al. 2009; Kim, et al. 2011; Smith, et al. 2011; Tian, et al. 2013; Park, et al. 2017; Zhao, et al. 2018) and the blind mole-rats (*Spalax galili*, BMRs) (Avivi, et al. 2001; Ashur-Fabian, et al. 2004; Shams, et al. 2004; Nasser, et al. 2005; Avivi, et al. 2010; Gorbunova, et al. 2012; Fang, Nevo, et al. 2014). NMRs and BMRs are phylogenetically distantly related subterranean rodents that evolved similar morphological and extraordinary physiological traits. To live in the burrow system, NMRs and BMRs evolved cylindrically shaped bodies, shortened limbs, strong claws, elongated incisors and degenerated visual system (Nevo 1979). They are highly tolerant to tissue hypoxia, resistant to oxidative stress induced by oxygen deficient and insensitive to acid induced by hypercapnia (Shams, et al. 2004; Kim, et al. 2011; Smith, et al. 2011; Fang, Nevo, et al. 2014; Park, et al. 2017). In addition, they are both extremely long-lived rodents that live more than 20 years and are able to suppress spontaneous and experimentally induced tumorigenesis (Seluanov, et al. 2009; Gorbunova, et al. 2012; Tian, et al. 2013), even though cases of cancer were reported recently in zoo-housed NMRs (Delaney, et al. 2016). They also evolved several species-specific traits, for example, eusociality (Jarvis 1981), poikilothermy (Buffenstein, et al. 2001) and lack of a circadian sleep rhythm in NMRs (Davis-Walton and Sherman 1994); enlarged brain structure related to sensing and orientation (Mann, et al. 1997; Kimchi and Terkel 2001), inconstant and changeful heart rate and increased muscle capillary density (Widmer, et al. 1997) in BMRs. These traits aroused general interests among researchers working on various filed such as evolutionary biology, genetics, aging and cancer research. Understanding the molecular bases of extraordinary traits in NMRs and BMRs may help us address some of the most challenging questions in biology and medicine, such as adaptations to extreme environments, mechanisms of hypoxia tolerance, mechanisms of aging and cancer resistance.

Genetic and genomic studies in NMRs and BMRs have uncovered the molecular bases of some extraordinary traits. For examples, genome sequencing revealed that the amino acid (AA) substitutions in NMR UCP1 protein are associated with its unique thermoregulation, and fast evolution of BMR embryonic haemoglobin gamma gene may contribute to hypoxia adaptation (Kim, et al. 2011; Fang, Nevo, et al. 2014). Comparative genomics has uncovered hundreds of positively selected genes (PSGs) and convergently evolved genes that may contribute to phenotypic evolution in NMRs and BMRs (Du, et al. 2015; Shao, et al. 2015; Davies, et al. 2018), including the convergence on the *SCN9A* gene (encoding Nav1.7), which leads to the repeatedly evolved acid insensitivity (Fang, Nevo, et al. 2014; Liu, Wang, et al. 2014). However, it should be noted that the positive selection and convergence rarely occurred in the same gene across subterranean linages and that evolutionary changes in different genes might produce similar phenotypes (Davies, et al. 2018). In addition, transcriptome analysis identified many differentially expressed genes that are associated with longevity, hypoxia and hypercapnia tolerance (Kim, et al. 2011; Malik, et al. 2011; Malik, et al. 2012; Fang, Nevo, et al. 2014; Fang, Seim, et al. 2014). For example, two p53 target genes, *CYTC* and *CASP9*, are downregulated in BMRs under hypoxia, which may contribute to cellular protection against hypoxia induced apoptosis (Fang, Nevo, et al. 2014). Also, *SMAD3* gene is upregulated in aging NMR brain, which may slow down the growth of cancer cells and contribute to cancer resistance (Kim, et al. 2011). Even though these studies identified many candidate loci that are potentially responsible for certain traits, functional assays are largely absent to validate their contributions. Moreover, most of these studies were focusing on the functional modifications of protein coding genes and many traits can still not be explained adequately. In this regard, to assess the contributions of other genetic makeups to subterranean adaptations is important and should not be ignored.

In addition to functional modifications of protein coding genes, other types of genetic changes can contribute to phenotypic evolution, such as pseudogenes (gene losses) (Albalat and Canestro 2016), regulatory element mutations (Wittkopp and Kalay 2012) and epigenetic modifications (Bonduriansky and Day 2009). Gene losses, which are usually associated with regressive evolution, were recently shown to play roles in adaptive phenotypic evolution more than we expected. Examples of adaptive gene losses include: i) in cetaceans, losses of *DSG4, DSC1, TGM5*, and *GSDMA* may contribute to the evolution of an epidermis morphology suit for aquatic environment and loss of *POLM* may contribute to the improved tolerance of oxidative DNA lesions (Sharma, et al. 2018; Huelsmann, et al. 2019); ii) in fruit bats, losses of *FAM3B* and *FFAR3* may help them adapt to sugar-rich diet by increasing insulin secretion and insulin sensitivity (Sharma, et al. 2018); iii) repeated losses of *PNLIPRP1* in herbivores may increase their capacity of fat storage, which is beneficial given that herbivores usually consume fat-poor diet (Hecker, et al. 2019). These observations are in consistence with the “less is more” hypothesis (Olson 1999). In terms of the subterranean mammals, however, almost all the studies addressing the role of gene losses in phenotypic evolution focused on eye degeneration (Kim, et al. 2011; Emerling and Springer 2014; Fang, Nevo, et al. 2014; Fang, Seim, et al. 2014; Prudent, et al. 2016; Emerling 2018), while the contribution of gene losses to subterranean adaptations has yet to be investigated.

In this study, to assess the contribution of gene losses to subterranean adaptations, we systematically identified gene losses in NMRs and BMRs, as well as their terrestrial relatives, guinea pigs and rats, which were used as terrestrial controls. Comparison of gene losses between subterranean and terrestrial lineages identified a batch of pseudogenes that might be associated with specific subterranean physiological traits including an altered DNA damage response and a specialized cardiovascular system. In addition, we provided functional evidence for the contribution of eliminating *TRIM17*, a gene independently lost from NMRs and BMRs, to hypoxia adaptation. Thus, our work provides new insights into the molecular mechanisms of subterranean adaptations of both NMRs and BMRs.

## Results and Discussion

### Compositions of the pseudogene repertoires

We first identified species-specific gene losses (unitary pseudogenes) in NMRs, BMRs, guinea pigs and rats by mapping a complete mouse protein set to their genomic sequences, followed by disablement (stop codons and frameshifts) detection and filtering by rigorous criteria (**supplementary fig. S1; see Materials and Methods**). The mouse genome was used as reference because mouse is currently the phylogenetically closest to NMRs and BMRs with high-quality genome assembly and annotations, which would mostly minimize the effects caused by phylogenetic distance. Although the difference in divergent time between each target species and mouse may affect the identified pseudogene numbers, the subsequent comparisons between subterranean and terrestrial control linages can help correct the phylogenetic bias as we focused on the proportion of functional similarity but not the absolute number of pseudogenes. Several factors affect the validity of detected gene losses including the misjudgement of orthologous sequences, genetic redundancy (Vavouri, et al. 2008; Qian, et al. 2010) and the false positives introduced by misjudgement of disruptive mutations, genome sequencing/assembly errors and annotation errors in reference genome. To address these issues and to obtain high-confidence gene losses for each species, we implemented a series of filtering steps (**supplementary fig. S1; see Materials and Methods**). Briefly, we first established the unitary status of each putative pseudogenic loci by i) removing loci hit by predicted proteins, intronless cDNA/expressed sequences or genes belong to large gene families such as olfactory receptors, zinc finger proteins and vomeronasal receptors; ii) removing genes with highly similar sequences at more than one locus in the genome; iii) removing genes without conserved genomic position between query species and mouse. Secondly, we removed false disruptive mutations generated by i) GeneWise alignment errors; ii) the sequencing and assembly errors; iii) potential annotation errors that resulted from the reference proteins with low transcript supporting level, or from downstream compensatory mutations that rescues the ORF disruption. As a result, 211, 211, 617 and 389 high-confidence unitary pseudogenes were identified from the NMRs, BMRs, guinea pigs and rats, respectively. To make our results more conservative, we further removed those pseudogenes without signatures of relaxed selection (**supplementary fig. S1; see Materials and Methods**) because gene dispensability and the following relaxed selective constraint is considered the hallmark of gene losses (Albalat and Canestro 2016). Finally, we obtained 167, 139, 341 and 112 gene losses in NMRs, BMRs, guinea pigs and rats, respectively (**fig. 1, table 1 and supplementary table S1**).

**Fig. 1.**
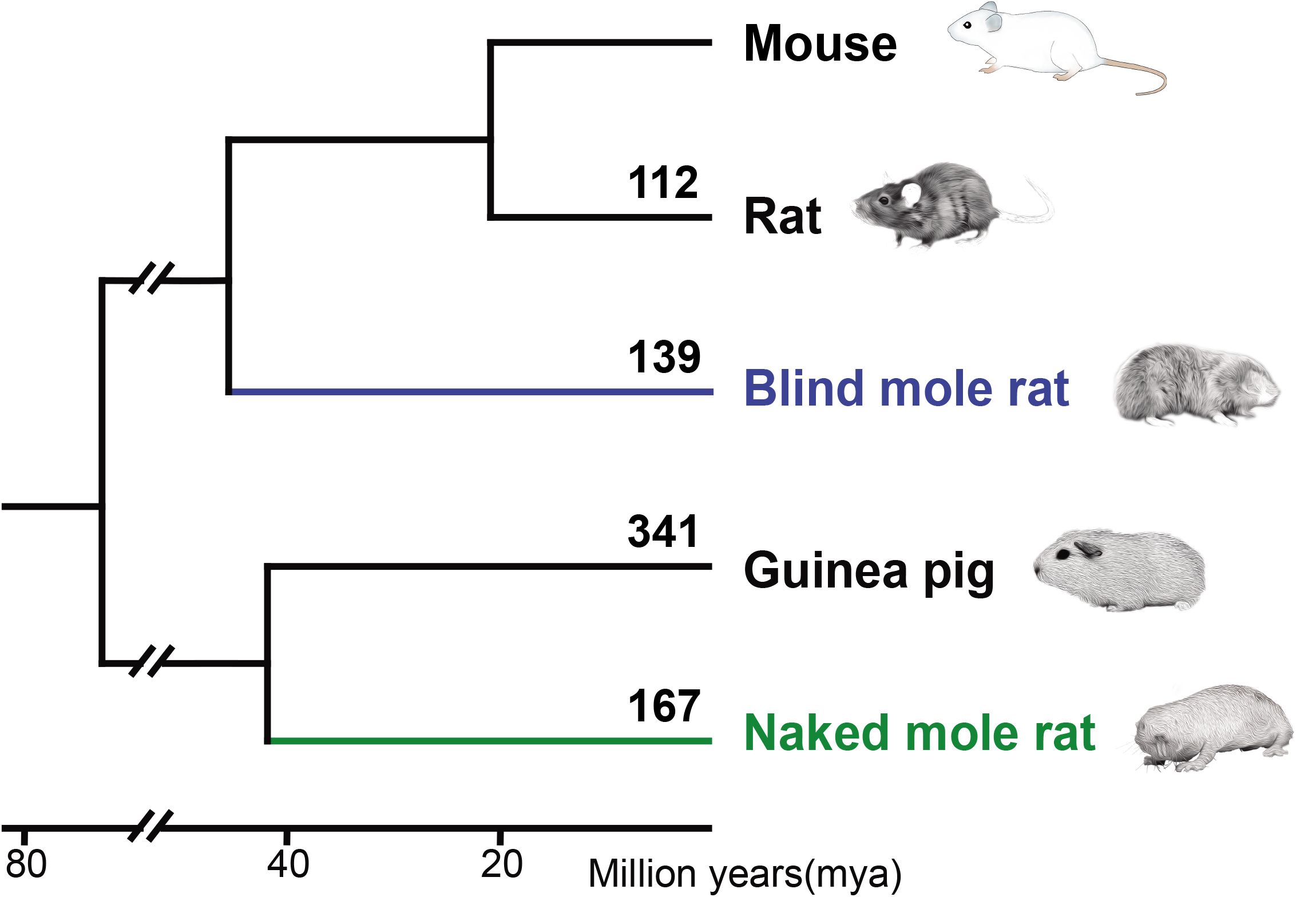
Gene losses identified in each species. The numbers of gene losses were labelled above each branch of the phylogenetic tree of species investigated.

**Table 1.** Statistics of each step of identifying gene loss events in NMRs, BMRs, guinea pigs and rats.

| Species | Overlapping best hits | Putative pseudogenic loci | Unitary pseudogenic loci | False positives removed | Selective constraints relaxed |
| --- | --- | --- | --- | --- | --- |
| NMRs | 14254 | 1362 | 883 | 211 | 167 |
| BMRs | 16546 | 1609 | 1023 | 211 | 139 |
| Guinean pigs | 15969 | 2695 | 2217 | 617* | 341 |
| Rats | 17396 | 2821 | 1701 | 389* | 112 |

\*Sequencing errors were not examined.

### Functional bias of gene losses in subterranean lineages

Next, we examined what kind of genes were preferentially lost in NMRs and BMRs, which are more likely to be associated with subterranean adaptations. Using mouse orthologs as surrogates, we firstly obtained all associated GO terms for all pseudogenes in each species. Functionally cross-related GO terms were clustered into functional groups and a subset of GO terms from each group that are overrepresented in subterranean lineage pseudogenes compared to subterranean lineage non-pseudogenes or control lineage pseudogenes (NMRs *vs*. guinea pigs and BMRs *vs*. rats) were identified as an enriched functional group (EFGs) (**see Materials and Methods**). Among the 24 clustered functional groups, a total of 10 EFGs were identified, with 5 in the NMR pseudogenes and 5 in the BMR pseudogenes (**fig. 2 and supplementary table S2**). For each EFG, the proportion of pseudogenes from the subterranean lineage is significantly higher than the pseudogenes from the corresponding control lineage and/or the functional genes from their own genome (**fig. 2 and supplementary table S2**). Genes in two EFGs might be associated with species-specific traits. For example, the EFG for “lipid metabolism” might be associated with the alternating membrane phospholipid composition in NMRs (Riccio and Goldman 2000; Mitchell, et al. 2007). The EFG “neural processes” might be associated with the enlarged brain volume in BMRs (Mann, et al. 1997; Kimchi and Terkel 2001). Four EFGs, namely, those for the “visual system”, “reproduction”, “DNA damage response” and “proteolysis”, were shared between the pseudogene lists from the two subterranean species. These repeatedly observed EFGs support a notion that subterranean adaptations are relatively restricted to where certain physiological process was recurrently modified to cope with stressful environments.

**Fig. 2.**
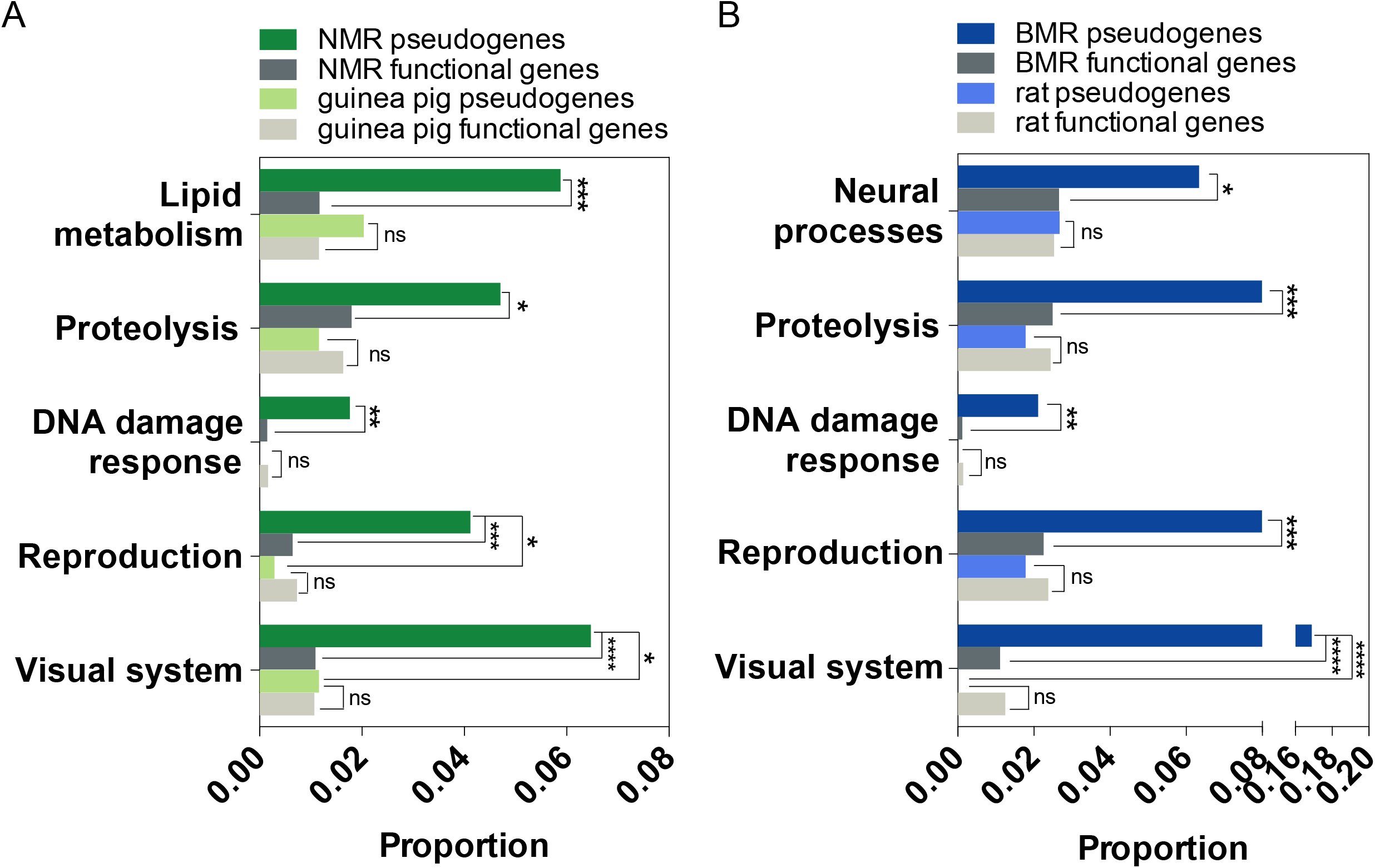
Functional groups enrichment analysis. Linage-specific and shared EFGs identified in NMR (**A**) and BMR (**B**) pseudogenes. The proportions of pseudogenes and functional genes background from each species in each EFG are shown and proportional difference in each EFG were tested with Fisher’s exact test. \**P* < 0.05, \*\**P* < 0.01, \*\*\**P* < 0.001, \*\*\*\**P* < 0.0001.

The EFG for “DNA damage response” is a good example to illustrate this notion. The highly hypoxic and hypercapnic burrow environments cause a significant oxidative stress on both BMRs and NMRs (Andziak, et al. 2006; Caballero, et al. 2006). The oxidative stress under hypoxic environment can cause DNA damages, especially DNA double-stranded breaks (DSBs) (Barzilai and Yamamoto 2004). We observed significant enrichments of NMR and BMR pseudogenes in “DNA damage response” EFG when compared to their own functional gene background (FDR corrected *P* = 0.0081 for NMRs and *P* = 0.0024 for BMRs, Fisher’s exact test) (**supplementary table S2**), while guinea pig and rat pseudogenes show no significance compared to their respective functional gene background (**supplementary table S2**). The genes in this EFG involve DNA damage checkpoint and DNA repair regulation, including *D7ERTD443E, NUDT16L1* and *EME2* in NMRs and *NEK11, D330045A20RIK* and *EME2* in BMRs (**fig 3A and supplementary table S2 and S3**). Among them, *D7ERTD443E* (*FATS*) and *NEK11* are checkpoint related genes that affect cell cycles and apoptosis by regulating p53/p21 and CDC25A, respectively (Melixetian, et al. 2009; Li, et al. 2010; Zhang, Zhang, et al. 2010; Yan, et al. 2014) (**fig. 3A**). Interestingly, both FATS and NEK11 are frequently deleted or downregulated in various human tumours (Zhang, Zhang, et al. 2010; Liu, Gao, et al. 2014), reflecting a beneficial effect of their functional losses on tumour cell survival, which is essential for tumor progression. This beneficial effect may probably be utilized by NMRs and BMRs to cope with hypoxic stress, particularly, prevent hypoxic cell death. The roles of FATS and NEK11 in tumour progression and hypoxia adaptation can be reconstituted by a *Spalax* p53 mutation, which mimics a human oncogenic mutation of p53 Lys-174 and was thought to be adaptive to hypoxia (Ashur-Fabian, et al. 2004). Supporting this idea, the NMR fibroblasts were shown to have a lower p21 response to γ-irradiation (IR), a reduced sensitivity to IR-induced senescence and resistance to IR-induced apoptosis (Zhao, et al. 2018).

**Fig. 3.**
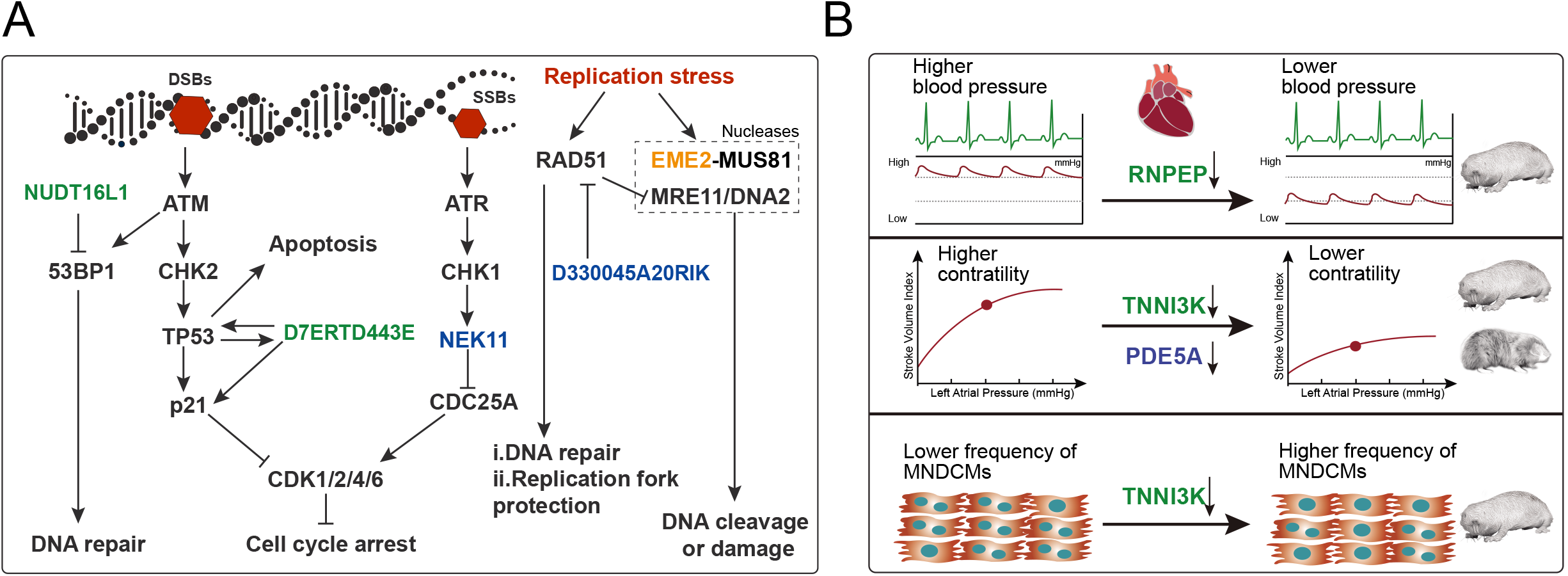
Functional convergences of gene losses between NMRs and BMRs in DDR pathway and cardiac features. **(A)** A part of DNA damage response pathway is represented. **(B)** Schematic diagrams of cardiac adaptations in NMRs and BMRs. Genes highlighted in green and blue are gene losses from NMRs and BMRs, respectively, and the gene highlighted in orange indicates shared gene loss between NMRs and BMRs.

DNA repair is another critical theme in DNA damage response and both NMRs and BMRs have evolved a higher DNA repair efficiency (Petruseva, et al. 2017; Domankevich, et al. 2018). Two negative DNA repair regulators, *NUDT16L1* (*TIRR*) and *D330045A20RIK* (*RADX*), were detected as pseudogenes in NMRs and BMRs, respectively (**supplementary table S2 and S3**). TIRR is a Tudor interacting repair regulator that inhibits the activity of 53BP1 and this inhibitory effect is utilized by cancer cells to increase genomic instability as *TIRR* locus was observed to be amplified in human cancers (Drane, et al. 2017). Depletion or silencing of TIRR markedly reduces the nuclear soluble levels of 53BP1, resulting in enhanced phosphorylation and association of 53BP1 with its effector proteins after induction of DNA damage (Drane, et al. 2017). In this regard, *TIRR* loss might help NMRs improve the DSBs repair efficiency by releasing more 53BP1 protein and enhancing 53BP1 activity in DNA repair. *RADX* encodes a single-strand DNA binding protein that prevents fork collapse by modulating RAD51 accumulation on replication forks (Dungrawala, et al. 2017). RAD51 is critical to reversal fork formation, reversed forks protection and homologous recombination repair (HR). However, either too much or too little of RAD51 accumulation on forks can cause DNA damage and fork collapse. RADX, acting like a buffer, competes with *RAD51* for binding to single-stranded DNA and ensures the right amount of RAD51 for reversal fork formation and protection (Bhat, et al. 2018). In fact, in normal cells, RADX inactivation leads to excessive RAD51 activity and DNA damages, despite RAD51 functioning as a DSB repair protein. In contrast, RADX inactivation in *BRCA2*-defìcient cancer cells, in which RAD51 function is reduced, is sufficient to rebalance RAD51 activity, protect forks from MRE11- and DNA2-dependent fork degradation (Dungrawala, et al. 2017; Bhat, et al. 2018). Due to the hypoxic environments, subterranean mammals suffer oxidative stress frequently, which causes replication stress and DNA lesions (Domankevich, et al. 2018) and prefer a higher RAD51 activity like *BRCA2*-defìcient cells. Therefore, RADX loss in BMRs might contribute to the enhanced DNA repair efficiency (Domankevich, et al. 2018) by rebalancing RAD51 functions, protecting against replication fork degradation and enhancing HR under a stressful condition. Interestingly, *EME2*, an endonuclease component gene that is capable of causing DNA damage under replication stress (Amangyeld, et al. 2014; Pepe and West 2014; Techer, et al. 2016), was independently lost from NMRs and BMRs (**fig. 3A and supplementary table S2 and S3**), which might help maintaining genome integrity by removing the potential source of DNA damages. Such observations suggest a contribution of these pseudogenes to the enhanced DNA repair efficiency in both species. Taken together, the similar direction of evolution of DNA damage response, i.e., attenuating apoptosis and enhancing DNA repair efficiency, indicates a functional convergence between NMRs and BMRs and highlights the importance of maintaining genome integrity under hypoxic conditions.

In addition to the shared EFGs, the pseudogenes in a functional group named “cardiovascular system” also show a functional convergence, even though no significant enrichments were observed (**fig. 3B and supplementary table S2**). This convergence might be associated with the low blood pressure in NMRs and low heart rate in BMRs, and both of them seem to be caused by changes in cardiac contraction (Arieli and Ar 1981; Widmer, et al. 1997; Grimes, Reddy, et al. 2014; Grimes, Voorhees, et al. 2014). In NMRs, *TNNI3K* and *RNPEP* were detected as pseudogenes (**supplementary table S3**). A recent study also reported that *TNNI3K* was pseudogenized in NMRs (Gan, et al. 2019). These two genes are involved in the contraction and action potential repolarization of cardiac muscle and blood pressure regulation. TNNI3K is a functional kinase and was shown to directly interact with cardiac troponin I (Zhao, et al. 2003). The overexpression of TNNI3K or its knockdown was shown to enhance or decrease the contraction of cardiomyocytes, respectively (Wang, Wang, Song, et al. 2013; Wang, Wang, Su, et al. 2013). In line with this, the natural knockout of TNNI3K in NMRs coincides with its low resting cardiac contractility (Grimes, Voorhees, et al. 2014) (**fig. 3B**). The other pseudogene, *RNPEP*, encodes Aminopeptidase B (Aurich-Costa, et al. 1997). Inhibition of RNPEP was shown to suppress the development of hypertension in spontaneously hypertensive rats (Aoyagi, et al. 1986), supporting its role in blood pressure regulation. Therefore, the abnormal regulation of blood pressure by losing *RNPEP*, together with low cardiac contractility by losing *TNNI3K*, might contribute to low blood pressure observed in NMRs (Grimes, Reddy, et al. 2014) (**fig. 3B**). In BMRs, *PDE5A*, which encodes a wide studied cGMP esterase, was detected as a pseudogene (**supplementary table S3**). PDE5A is involved in various cardiomyopathies including cardiac hypertrophy and myocardial infarction (Takimoto, et al. 2005; Perez, et al. 2007; Tedford, et al. 2008; Giannetta, et al. 2012). Interestingly, like TNNI3K, the inhibition of PDE5A was also shown to associate with suppressed contractility (Borlaug, et al. 2005) (**fig. 3B**). Even though the cardiomyocyte contractility in BMRs has yet to be determined, the heart rate of BMRs was observed to be lower than rat under both normoxia and hypoxia conditions (Arieli and Ar 1981), which might be caused by lowered cardiac contractility. Interestingly, inhibition of *TNNI3K* or PDE5A was shown to be cardioprotective against several cardiomyopathies (Takimoto, et al. 2005; Perez, et al. 2007; Tedford, et al. 2008; Giannetta, et al. 2012; Vagnozzi, et al. 2013). Moreover, TNNI3K inhibition was shown to increase the frequency of the mononuclear diploid cardiomyocyte (MNDCM) population and to elevate cardiomyocyte proliferation after injury (Patterson, et al. 2017; Gan, et al. 2019) (**fig. 3B**). In this regard, the cardioprotective effects conferred by *TNNI3K* or *PDE5A* losses in each species are likely to be adaptive because cardiomyocytes are vulnerable to oxidative stress, which poses a challenge to cardiomyocyte survival (Gottlieb, et al. 1994; Yellon and Hausenloy 2007).

### Potential roles of convergent gene losses (CGLs) in subterranean adaptations

It has been shown recently that CGLs contribute to repeatedly evolved adaptations (Meyer, et al. 2018; Sharma, et al. 2018; Hecker, et al. 2019). To reveal subterranean adaptations that were contributed by CGLs, we compared the pseudogene lists from NMRs and BMRs. 20 CGLs were observed between NMRs and BMRs (**supplementary table S3 and S4**). The hypergeometric test shows that these CGLs significantly outnumbered the random expectation (1.42, *P* = 1.25×10^-17^, **fig. 4B**). To examine the extent to which the number of CGLs are affected by similarity of ecological niches, we further paired the four species according to the similarities and differences in their living environments. Among 4 pairs of species, NMRs-BMRs (N-B) and guinea pigs-rats (G-R) live in similar environment (homotypic pair), while NMRs-rats (N-R) and BMRs-guinea pigs (B-G) live in different environment (heterotypic pair). Comparison of pseudogene lists between paired species shows that homotypic N-B pair is the most prominent with highest number of CGLs (**fig. 4A**). In contrast, the number of CGLs from homotypic G-R pair is significantly lower than N-B pair (5/453 vs. 20/306, *P* <0.0001, Fisher’s exact test) (**fig. 4A**). The CGLs number from the homotypic N-B pair is significantly higher than that from heterotypic B-G pair (20/306 vs. 7/480, *P* = 0.0002), while there is no significant difference between the CGLs number from the homotypic G-R pair and that from the heterotypic N-R pair (**fig. 4A**). These results suggest that the CGLs between NMRs and BMRs might result from the similar response to lifestyle transition from aboveground to underground and probably contribute to the evolution of their shared traits.

**Fig. 4.**
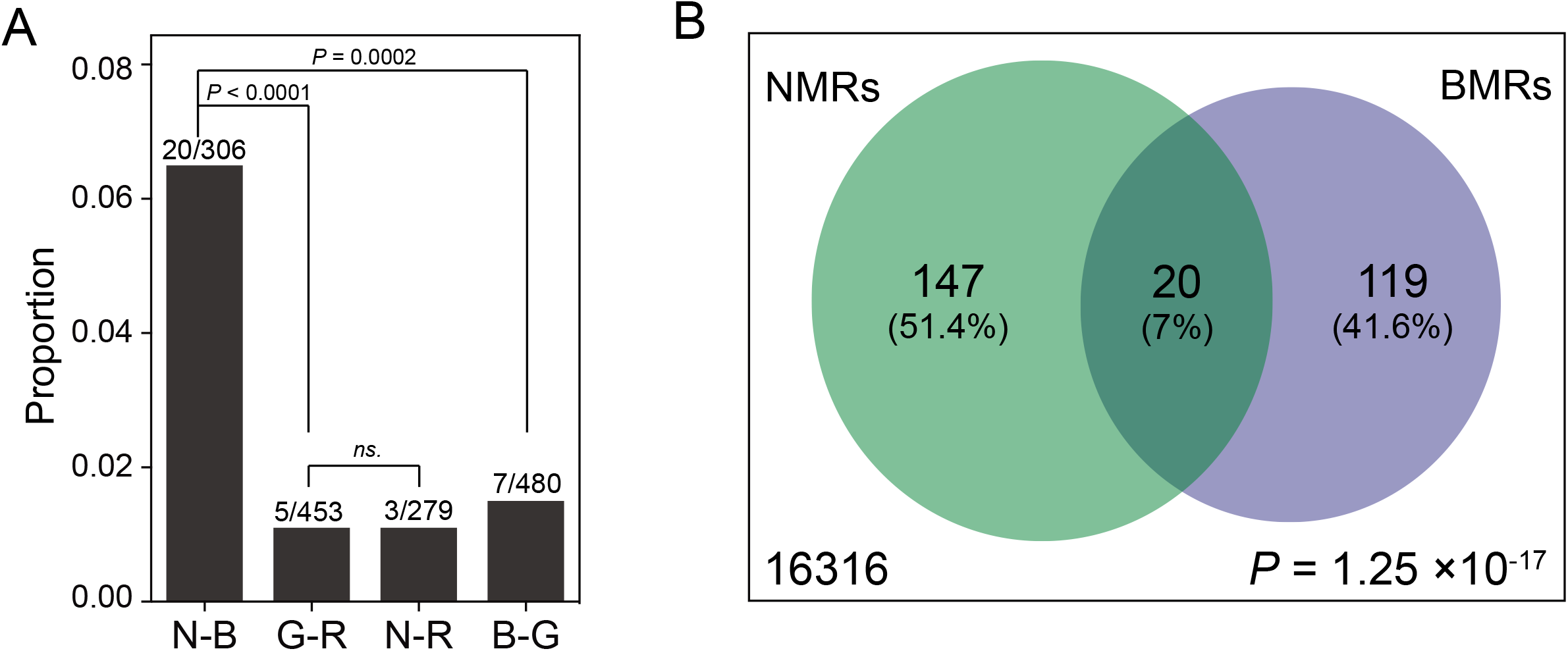
Convergent gene losses in species pairs from similar and distinct environments. **(A)** Proportional difference of overlapping pseudogenes among 4 species pairs including NMRs-BMRs (N-B), guinea pigs-rats (G-R), NMRs-rats (N-R) and BMRs-guinea pigs (B-G). The numbers of overlapping and total pseudogene from each pair are labelled above the bars. *P*-values are from two-tailed Fisher’s exact test. **(B)** Venn diagram showing the one-to-one orthologous genes (large box) and overlapping pseudogenes in NMRs-BMRs pair. *P*-value from a hypergeometric test indicates that gene losses in NMRs and BMRs are not independent from each other.

To associate overlapping pseudogenes with subterranean traits, we leveraged knockout mice phenotypic information (MPI) from literatures or MGI database and genetic associations from the public NHGRI-EBI GWAS catalog (MacArthur, et al. 2017). Degenerated visual capacity is one of the most significant traits that shared between NMRs and BMRs. By examining the MPI, we found that 5 convergent gene losses, including *GUCY2F, ABCB5, RP1L1, CRB1* and *ARR3* (Deming, et al. 2015), are vision related and knockout mice of each gene show visual abnormality (**supplementary table S5**). It is worth noting that we also identified many other vision related genes that lost specifically in each species, including genes reported by previously studies (Kim, et al. 2011; Emerling and Springer 2014; Fang, Nevo, et al. 2014; Prudent, et al. 2016; Emerling 2018; Sharma, et al. 2018), and new vision related pseudogenes, such as *CRB1* in BMRs **(supplementary table S6**). Thus, these gene losses collectively contribute to the eye degeneration in NMRs and BMRs. We also searched the significant genetic associations (*P* < 1e-06) for all overlapping genes that curated in the NHGRI-EBI GWAS catalog. Interestingly, we found that variants in 5 genes, including *ABCB5, LETM2, INMT, RP1L1* and *CRB1*, are significantly associated with cardiovascular traits including PR interval, pulse pressure and systolic blood pressure (Deng, et al. 2013; Evangelou, et al. 2018; Feitosa, et al. 2018; Ramírez, et al. 2018; Giri, et al. 2019)(**supplementary table S7**). Among them, except that SNP in *INMT* gene is located in coding region, all of the rest variants are located in regulatory regions or introns. Even though further investigations are needed, these data imply the contributions of losing such genes to cardiovascular adaptations in both NMRs and BMRs, especially blood pressure control under hypoxia. More importantly, we noticed that *ABCB5*, *RP1L1* and *CRB1* are also vision related pseudogenes. Therefore, these genes may control multiple traits, in this case, the degenerated eyes and cardiovascular traits, simultaneously. This phenomenon is also observed in other vision related genes. For example, knockout mice of *BFSP1*, a BMRs vision related pseudogene, show both visual abnormalities and enlarged heart (MP:0000274). The phenotypic effects outside the visual system are indicative of gene pleiotropy and arise a question of why NMRs and BMRs lost these genes despite their non-visual functions. One possibility is that the non-visual phenotypic effects generated by losing such visual genes provides survival benefits because they affect cardiac features, which is important for hypoxia adaptation. If this is the case, some vision related gene losses should not simply be interpreted as the consequences of relaxed selection, rather, they probably have played positive roles in subterranean adaptations. A recent study in human populations demonstrated that many myopia-related mutant alleles are associated with reproductive benefits, and the reproduction-related selection probably contributes to the myopia epidemic inadvertently (Long and Zhang 2020).This is well supporting our above hypothesis. When further investigations are conducted and more data are accumulated in the future, we expect to have a better understanding of this issue.

The functional studies also provide clues regarding the potential roles of gene losses in subterranean adaptations. For example, as mentioned earlier, loss of *EME2* could probably alleviate DNA damages induced by oxidative stress under hypoxic condition (Amangyeld, et al. 2014; Pepe and West 2014; Techer, et al. 2016). Another identified pseudogene, *TRIM17*, is also quite intriguing. It has been suggested that TRIM17 is involved in trophic factor withdrawal-induced neuronal apoptosis (Lassot, et al. 2010). Considering that both NMRs and BMRs are highly tolerant to hypoxia induced brain injury (Avivi, et al. 2010; Park, et al. 2017), it is possible that independent loss of *TRIM17* in NMRs and BMRs contribute to protecting neural cells against hypoxia induced apoptosis. However, no direct links have been established between hypoxic stress and the actions of TRIM17, neither *in vivo* nor *in vitro*. It is also possible that other uncharacterized functions of TRIM17 are associated with some other subterranean traits. For example, *TRIM17* knockout mice show physiological traits that are not involved in nerve system, including increased circulating alkaline phosphatase level (MP: 0002968) and increased circulating thyroxine level (MP: 0005477) (**supplementary table S5**). Therefore, functional investigations for these less well-characterized genes would provide insights into the roles of their functional losses in subterranean adaptations.

### Functional loss of *TRIM17* contributes to neuroprotection under hypoxia *in vitro*

We then chose *TRIM17* to perform the functional survey to check whether loss of *TRIM17* is involved in protecting neural cells against hypoxia-induced apoptosis, as preventing cell death and injury, especially neuronal cells, is a critical theme in adapting to hypoxic environments (Ramirez, et al. 2007). First, we examined the pseudogenization status of *TRIM17* and their possible correlation with hypoxia. Disruptive mutations were confirmed by manual inspection of the alignments, showing that *TRIM17* is inactivated by one premature stop codon in NMRs and 1 stop codon/4 indels in BMRs (**fig. 5A**). The RING finger domain of TRIM17 is used to execute its E3 ligase function (Lassot, et al. 2010), however, it is disrupted in both species. We also examined the frequency of disruptive mutations in 11 BMR populations (Li, et al. 2015), and no segregations were found (Data not shown). In NMRs, the premature stop codon was also verified by the available transcriptomic reads and genomic sequence of another individual (**supplementary table S3**). Furthermore, *TRIM17* shows signature of relaxed selective constraint in both species, even though not significant (NMRs: *k* = 0.69, *P* = 0.1338; BMRs: *k* = 0.72, *P* = 0.1214) (**supplementary table S1**). These results together confirmed the elimination of an intact *TRIM17* in both NMRs and BMRs. Interestingly, we noticed that *TRIM17* was independently lost at least two times in cetaceans, with a shared 1bp deletion in toothed whales (Monodontidae, Phocoenidae and Delphinidae) (Sharma, et al. 2018) (**fig. 5B**) and a shared premature stop codon in baleen whales (Balaenopteridae) (**fig. 5C**). Inactivating mutations were also detected in several species in other families including right whale (*Eubalaena japonica*), pygmy sperm whale(*Kogia breviceps*) and boutu (*Inia geoffrensis*) (Data not shown). Moreover, *TRIM17* was also lost in two pangolins (*Manis pentadactyla* and *Manis javanica*) due to a shared premature stop codon (**fig. 5D**) (Sharma, et al. 2018). Like NMRs and BMRs, cetacean and pangolins experience hypoxic stress during their lifetime (Weber, et al. 1986; Ramirez, et al. 2007; Choo, et al. 2016). Therefore, independently losses of *TRIM17* in distinct hypoxia tolerant taxa probably suggest a relevance of *TRIM17* losses in hypoxia adaptation.

**Fig. 5.**
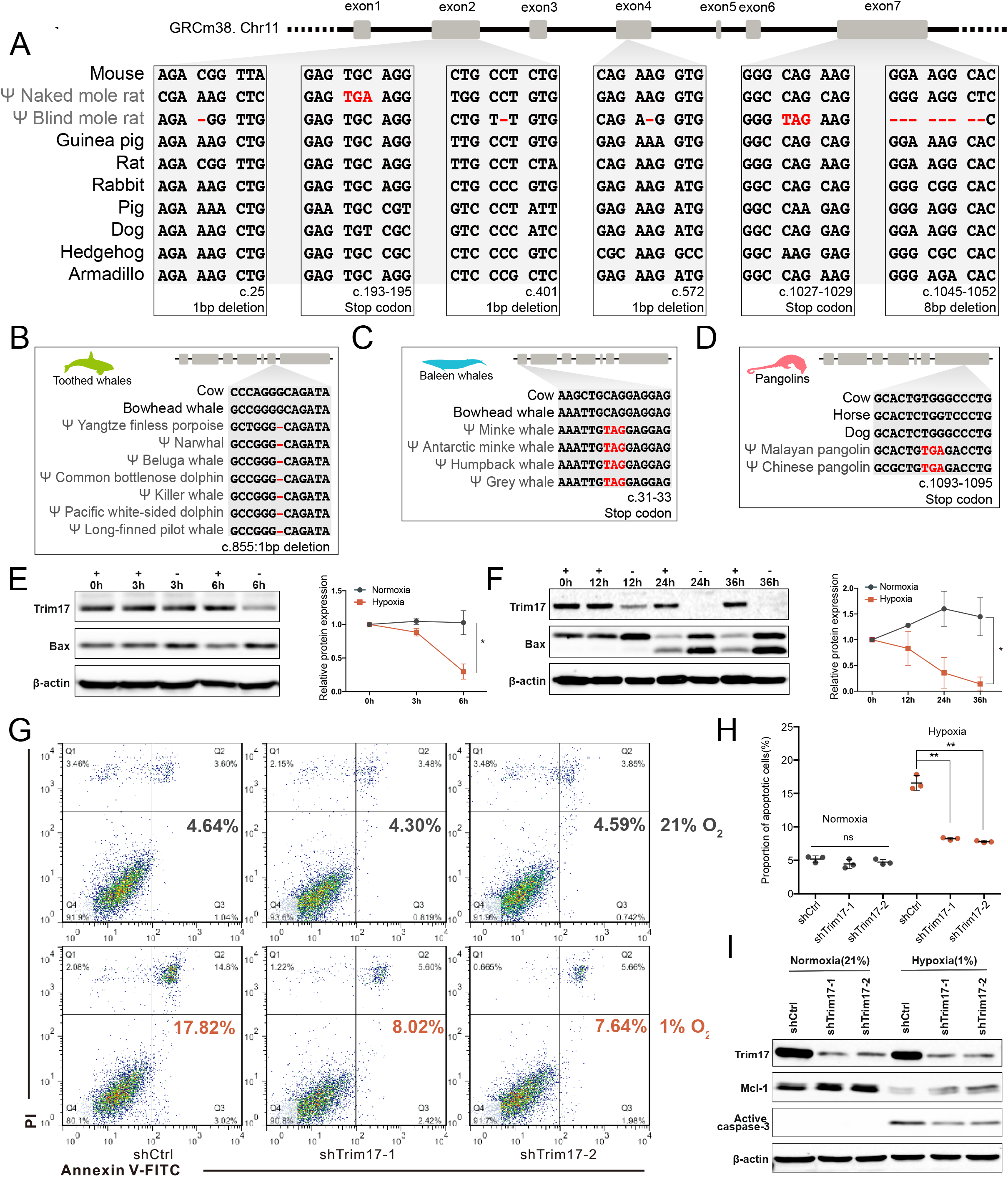
Functional tests of *TRIM17* for its role in hypoxia adaptation. **(A)** Coding sequence alignment of *TRIM17* orthologous genes from mouse, NMR, BMR and 7 other mammals. One stop codon in NMRs and one stop codon/4 frameshifts in BMRs were observed that resulted in disrupted coding potential of *TRIM17* gene. (**B** to **D**) Examples of inactivating mutations in toothed whales (B), baleen whales (C) and pangolins (D). The positions of all mutations shown in this figure are labelled under the corresponding alignment using mouse Trim17 coding sequence as reference. (**E** to **F**) Immunoblot analyses for TRIM17 protein level changes during hypoxia treatments in primary cortical neurons (**E**) and N2a cell lines (**F**). “+” indicates hypoxia treatment and “-” indicates normoxia treatment. Quantification and relative foldchange analysis of protein level showed a significantly decrease of Trim17 protein induced by hypoxia in both primary cortical neurons and N2a cells. Data from three independent assays were used to perform paired two-tailed t-test, \**P* < 0.05. (**G** to **I**) Knockdown of Trim17 expression by shRNAs (shTRIM17-1 and shTRIM17-2) provides neuroprotection under hypoxia (1% O_2_) compared to control (shCtrl). Apoptotic cells are detected by staining cells with PI and Annexin V-FITC followed by flowmetry analysis. Cells with Trim17 knockdown showed significantly lower apoptosis rate under hypoxia compared to control, while no significant changes were observed under nomorxia (**G, H**), paired two-tail t-test, \**P* < 0.05, \*\**P* < 0.01. Immunoblot analysis for corresponding cells shows elevated Mcl-1 protein level and decreased activated caspase-3 protein level under hypoxia in cells with Trim17 knockdown (**I**).

To illustrate this idea, we next examined the changes in Trim17 expression in response to hypoxia in both Neuro-2a (N2a) cells and mouse primary cortical neurons. An immunoblot analysis shows that in contrast to trophic factor withdrawal, the Trim17 protein level is significantly decreased in both N2a cells and primary neurons after treating with hypoxic stress (1% O_2_) (**fig. 5E-F**), suggesting a hypoxia-responsive regulatory mechanism for Trim17 protein. We further tested whether Trim17 knock-down confers a neuroprotective effect under hypoxia. N2a cell lines stably expressing shRNAs that targeted Trim17 and control shRNA were constructed and treated under hypoxic condition (1% O_2_) for 48 h, followed by an apoptosis analysis via flow cytometry. In contrast to the increased apoptotic rate in the control cells (shCtrl), the Trim17 knock-down cells (shTrim17-1,2) showed significantly reduced apoptosis under hypoxia compared with normoxia (21% O_2_) (**fig. 5G-H**). As Trim17 was shown to initiate apoptosis through the protostome-dependent degradation of Mcl-1 (Magiera, et al. 2013), we examined whether this is also the case under hypoxia. An immunoblot analysis revealed elevated Mcl1 protein level and consequently decreased active caspase-3 protein under hypoxia when compared to the control (**fig. 5I**). Taken together, these results suggest that by preventing MCL-1 protein from degradation, eliminating TRIM17 function confers neuroprotection against hypoxia, and therefore, probably contributes to hypoxia tolerance. In the future, further investigations addressing the linkages among the components of phenotype-genotype-fitness continuum would provide us a better understanding of gene loss and adaptation.

### Gene losses and PSGs cooperatively contribute to subterranean adaptations

Physiological adaptations to hypoxic underground niches, such as modifications for DNA damage response (DDR) and cardiovascular system, are usually complex and expected to be involving multiple genes. We noticed that many other genomic studies had uncovered several PSGs that are also involved in DDR and heart functions in NMRs and BMRs (Kim, et al. 2011; Fang, Nevo, et al. 2014; Fang, Seim, et al. 2014; Davies, et al. 2018). Unlike gene losses, PSGs show a signature of selection that can be easily detected, and thus could be assigned as adaptive. To examine whether PSGs also show enrichments in functional groups that similar to EFGs in pseudogenes, we collected PSGs from previous genomic studies and compiled a PSG set for NMRs (Kim, et al. 2011; Fang, Seim, et al. 2014) and BMRs (Fang, Nevo, et al. 2014; Davies, et al. 2018), respectively (**see Materials and Methods**). Interestingly, for some of the EFGs in pseudogenes, we also observed similar functional enrichments in their respective PSG sets, including functional groups related to “DNA damage response”, “lipid metabolism” and “neural processes” (**supplementary table S8**). Moreover, consistent with the phenotypic convergence in heart contractile function, PSGs in functional group of “cardiovascular system”, for which no enrichment was observed in pseudogenes, also show significant enrichment (**supplementary table S8**). In fact, several genes involving cardiac muscle contraction in both species were under positive selection. These observations highlight the collaborative contributions of not only different genes, but also different types of genetic makeups to physiological adaptations. For example, proteinprotein interaction analysis (STRING) revealed that in both NMRs and BMRs, interaction networks involving DNA damage response contain both pseudogenes and PSGs, and both of them interact with other critical mediators such as *ATM, ATR* or *TP53* (**fig. 6**). Such simultaneous gain-of-function and lost-of-function changes in multiple genes among a network reflect the strong rewiring processes in response to drastic environmental changes, and rise a question of how gene losses and PSGs work together to achieve a functional benefit.

**Fig. 6.**
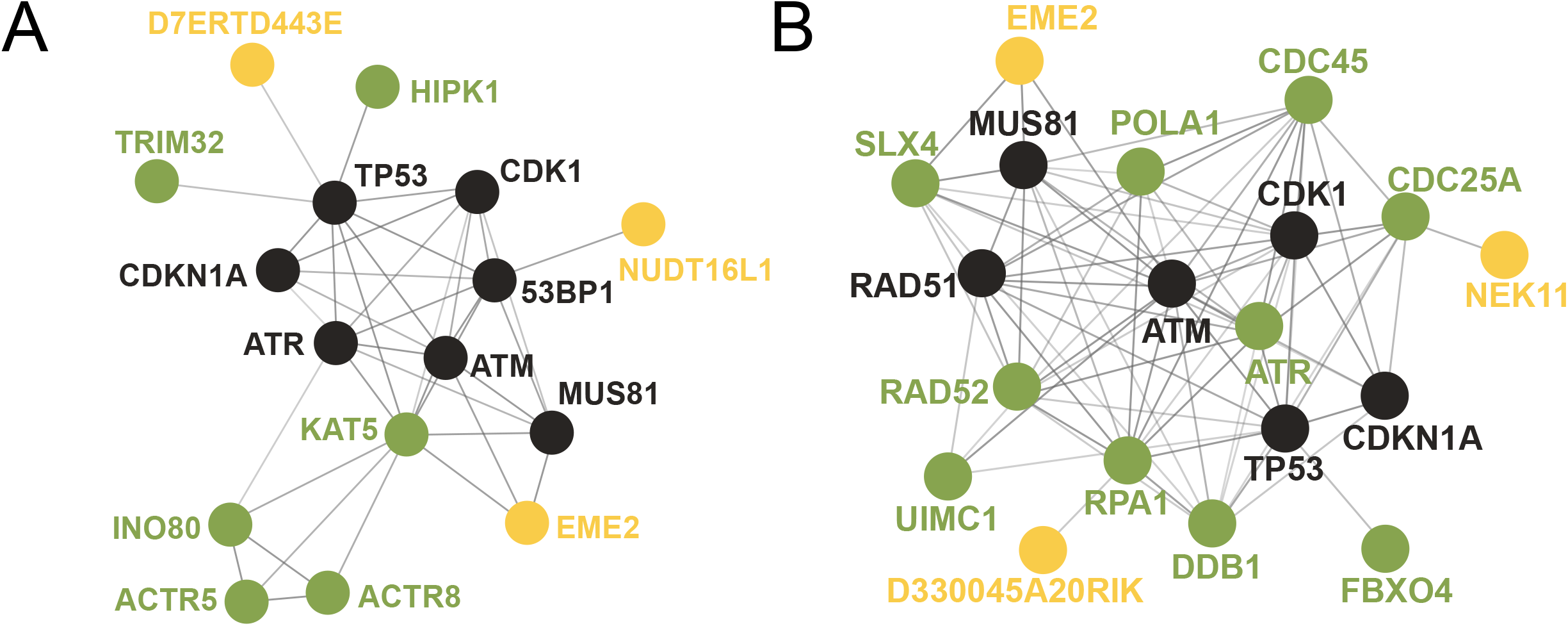
Interacting network of DDR related proteins. Different genes with different types of genetic changes are interacting with each other in both NMRs (A) and BMRs (B). PSGs are highlighted in green and gene losses are highlighted in yellow, and core mediators of DDR are highlighted in black. Interaction information were derived from STRING database (https://string-db.org/, Version 11.0).

One can easily speculate that gene losses and PSGs may have similar functional effects that together result in an additive effect on a trait. For example, in the case of NMR DDR network, two genes, a PSG *TRTM32* and a pseudogene *D7ERTD443E* (*FATS*) interacts directly with *TP53* (**fig. 6A**). Both of them are ubiquitin ligase, but they regulate protein stability of TP53 in opposite direction, with a ubiquitination-mediated stabilization by FATS and a ubiquitination-mediated degradation by TRIM32 (Liu, Zhang, et al. 2014; Yan, et al. 2014). Pseudogenization of *FATS* eliminates one layer of positive control of TP53 protein stability and alternatively, facilitates negative regulation together with TRIM32, or other negative regulators (Vousden and Lane 2007). Another possibility considers the “sign-epistatic” interaction: either types of genetic change could be initially deleterious *per se*, while before its elimination by purifying selection, one of them may serve as the stepping stone for another one to reach the otherwise inaccessible complex, beneficial functions by fitness valleys crossings (Lenski, et al. 2003; Weinreich, et al. 2005; Cowperthwaite, et al. 2006; Weinreich, et al. 2006; Covert, et al. 2013). It is also possible that losing genes serves as a compensation strategy to buffer the deleterious effects of beneficial mutations in highly pleiotropic genes or *vice versa*, and such compensations were recently observed experimentally in a genome-wide scale study in baker’s yeast (Szamecz, et al. 2014). It is more likely that all these possible interactions exert their phenotypic effects simultaneously, due to the rewiring of a complex genetic network underlying physiological adaptations, especially to hypoxia, must have been strenuous. The roles of different types of genetic changes in complex adaptations in mammals have rarely been discussed. Our study implies the caveats of exploring the genetic mechanism of an adaptation by only looking at a single type of genetic change, e.g., the PSGs. Rather, a systematic perspective with details regarding the division of labour for different types of genetic makeups would be preferable.

### Conclusions

With combination of comparative genomic approaches and *in vitro* experimental validations, our study, for the first time, uncovered that gene losses might contribute to the evolution of specialized physiological traits in both NMRs and BMRs, including altered DNA damage response, specialized cardiovascular system and neuroprotection. We also highlighted that the genetic mechanisms underlying physiological adaptations to subterranean niches are complex, involving not only multiple genes but also different types of genetic makeups. Taken together, our study provides new insights into the molecular underpinnings of subterranean adaptations, highlighting the important roles of gene losses. Several pseudogenes identified in BMRs and NMRs might have implications for medical researches involving aging and longevity, neuroprotection in ischemic/hypoxic injury and cardiovascular diseases. Moreover, our study suggests an integration of different genetic makeups in future studies and also has implications for the comprehensive understanding of subterranean adaptation.

## Materials and Methods

### Genomic data

The genome sequences of each species analysed in this study were retrieved from the Ensembl database (http://asia.ensembl.org/) (mouse: GRCm38.p2, release 75; rat: Rnor_6.0; guinea pig: Cavpor 3.0, release 89) and NCBI database (https://www.ncbi.nlm.nih.gov/) (NMR: HetGla_female_1.0; BMR: S.galili_v1.0) together with its annotated CDS and protein sequences. The CDS and protein sequences from rabbit, which were used to estimate the evolutionary rate, were retrieved from the Ensembl database (OryCun2.0).

### Identification of gene losses

Basically, the gene losses in each species are defined as genes harbouring ORF-disrupting mutations, including premature stop codons and/or frameshifts with intact 1:1 orthologs in the mouse genome. Orthologous relationships were first established by mapping the longest protein sequence for each mouse gene to each query species by simultaneously using BLAT (Kent 2002) and genBlastA (She, et al. 2009). The genomic regions identified by both BLAT and genBlastA were considered as orthologous sequences and were extracted with 5000 bp downstream and upstream flanking sequences to predict the gene structure and ORF by GeneWise (Birney, et al. 2004). Alignments containing premature stop codons and/or frameshifts were identified, and the corresponding genomic regions were defined as putative pseudogenic loci. To establish the unitary status of each putative pseudogenic locus, several stringent filtering steps were executed as follows: 1) Loci hit by proteins belonging to large gene families including olfactory receptors, zinc finger protein and vomeronasal receptors were removed because of their high sequence similarity. 2) Loci hit by predicted proteins or intronless cDNA/expressed sequences were removed because they are unlikely to be real or unitary (Zhang, Frankish, et al. 2010). 3) To remove the potential functional redundancy, e.g., gene duplication or retrotransposition, BLAT mapping hits from the initial step were reanalysed to remove loci with homologous sequences across the genome. In detail, for each locus, we firstly removed hits with E-value higher than 1e-03, and then we assembled the remaining hits into “copies” of this pseudogene along the mouse protein coordinates. Each copy includes non-overlapping hits that together cover the mouse protein as long as possible. Simply, for each “copy” with total protein length longer than 80% of the corresponding mouse protein, we consider it as a potential functional redundancy and remove it. 4) We examined the conserved genomic position for each locus to ensure the orthologous relationship. Pairwise genomic alignments between mouse and each query species were downloaded from Ensembl database. For each locus, the genomic coordinates identified by us was compared to the coordinates inferred from a pairwise genomic alignment. Only pseudogenic loci with conserved genomic position were retained for further consideration. Furthermore, for genes discussed in the main text, we also manually examined the conserved gene order according to mouse genome annotation in Ensembl. To generate high-confidence pseudogene sets, false positives with disruptive mutations introduced by GeneWise, sequencing errors or annotation errors were removed through the following steps: a) in-house Perl script filtering in combination with manual inspection was performed to remove false positives introduced by GeneWise. Pseudogenes with a single disruptive mutation located at the end of the coding region (which resulted in truncated sequences longer than 90% of predicted intact protein (MacArthur 2012)), at the intron and exon boundaries or in regions with low similarity in GeneWise alignments were removed. Pseudogenes with multiple disruptions were manually inspected to remove short or low-quality GeneWise alignments. For the genes discussed in the main text, we realigned the mutated exons on mouse genome sequence using CESAR (Sharma, et al. 2017) to remove spurious disruptive mutations caused by evolutionary splice site shifts. b) to remove the sequencing and assembly errors, we retrieved the raw sequencing reads from NMRs and BMRs to check the validity of each disruption in the corresponding reads. For genes discussed in the main text, we further checked their validity using additional genomic/transcriptomic resources, i.e. raw transcriptomic reads of different tissues (Kim, et al. 2011) and genomic sequences of a male individual for NMRs (HetGla_1.0); resequencing reads of 11 individuals for BMRs (Li, et al. 2015). c) the coding potential of the corresponding genes was examined by checking the Ensembl transcriptional support level (TSL, level 1/2) and CCDS assignment, and those satisfying any one of the criteria were retained. d) pseudogenes with only two compensatory frameshifts were manually examined and removed because it provided little evidence for gene loss.

### Detection of relaxed selection

Genes with disruptive mutations on coding sequence do not necessarily indicate gene dispensability. Such disruptive mutations might be rare genetic variation or sequencing/assembly errors. To address this issue, we detected signature of relaxed selection for each pseudogene by RELAX (Wertheim, et al. 2015) and using a phylogenetic tree with rabbits as the outgroup. In detail, the coding sequences (CDS) and their translations (with disruptive mutations removed) of each pseudogene were extracted from the GeneWise alignments. The protein sequences were used to identify 1:1 orthologous gene in the rest of the species for each pseudogene by reciprocal best BLAST hit (RBBH) method, and then, the results were clustered into orthologous groups. For each orthologous group, CDS were used to construct codon-based multiple sequence alignment with a phylogeny-aware alignment algorithm as implemented in the PRANK program (Loytynoja and Goldman 2005). To obtain reliable CDS alignments, we set up rigorous filtering criteria including: i) removing gaps and ambiguous bases together with 6 bp flanking sequences with Gblocks (Castresana 2000) using the default parameters; ii) removing alignment fragments with low quality (Gblocks, default parameters); and iii) removing alignments with lengths of less than 150 bp. Alignments containing at least two sequences were assigned to a specific tree file. In each alignment, the species in which this gene loss occurred was set as “Foreground” branch and the rest of species were set as “Background” branch. Only pseudogenes with a higher dN/dS in foreground and a selection intensity parameter *k* < 1 (k > 1: intensified selection; k < 1: relaxed selection) were retained.

### Gene ontology

Orthologous mouse genes were used as surrogates to infer the functions of the pseudogenes or PSGs in corresponding species. Only pseudogenes and PSGs with orthologous relationship with mouse (established by BLAT and genBlastA mapping) were further considered. To identify the EFGs from pseudogenes, the associated GO terms for each gene were extracted from go.obo (released February 20, 2018), and the mouse gene annotation files (*Mus musculus*, released September 29, 2017) were downloaded from the Gene Ontology Consortium (http://geneontology.org/). NMRs and BMRs genes with GO terms involved in certain biological processes and systems were manually clustered into functional groups based on the keywords. For each functional group, we then compared the proportional differences of genes from subterranean lineage pseudogene lists, subterranean lineage non-pseudogene lists (with one2one mouse orthologs), control lineage pseudogene lists and control lineage non-pseudogene lists (with one2one mouse orthologs). For example, A subset of GO terms with a significantly higher proportion of NMR pseudogenes compared to that of NMR non-pseudogenes and/or that of guinea pig pseudogenes was defined as an EFG. For PSGs, EFGs were defined only when significantly higher proportion of NMR (or BMR) PSGs in a subset of GO terms was observed compared to that of NMR (or BMR) non-PSG background (with one2one mouse orthologs). Fisher’s exact test was used to test the significances of differences in proportion and FDR-adjusted *P*-values for each multiple test were reported using p.adjust() function in R.

### Cell culture

Primary cortical neurons were prepared from E16-E18 embryos of Kunming mice. Specifically, embryonic cortices were dissected in cold DMEM (Gibco) supplemented with 10% fetal bovine serum (Gibco) and then digested using 2 mg/ml papain (CHI). The resulting cell suspension was centrifuged at 1500 rpm for 5 mins. The cells were gently resuspended in DMEM supplemented with 10% FBS and gentamycin (50 μg/ml, Solarbio) and filtered through a 70 μm cell strainer (Corning Falcon). Then, the cells were seeded at a density of 1.2 × 10^6^ cells/ml in culture dishes coated with 20 μg/ml poly-D-lysine (Sigma). The neurons were cultured at 37°C in a humidified incubator with 5% CO_2_/95% air, and the medium was replaced 6 hours afterwards with neurobasal medium (Gibco) supplemented with B27 supplements (Gibco) and L-glutamine (Gibco). After 6 days *in vitro* (DIV 6), the neural cultures were subjected to hypoxia treatment and protein extraction. N2a cells was purchased from the Cell Bank of Type Culture Collection of the Chinese Academy of Sciences (Shanghai; Institution Code: CBTCCCAS) and were cultured at 37°C in a humidified incubator with 5% CO_2_/95% air with MEM (Gibco) supplemented with 1.5 mg/ml NaHCO3, 0.11 mg/ml sodium pyruvate (Gibco), and 10% FBS.

### Hypoxia treatment paradigms

The hypoxia (1% oxygen concentration) for the cell culture treatment was achieved using a humidified hypoxia chamber (Proox, model C21, BioSpherix) set at 37°C, 1% O_2_, and 5% CO_2_ and maintained with a N_2_ supply. The primary cortical neuron cultures and N2a cells (including the stable cell lines) were treated at 1% hypoxia for various periods of time. As controls, cells were cultured at 37°C in a humidified incubator with 5% CO_2_ and 95% air.

### Immunoblot analysis

Primary cortical neuron and N2a cells (including stable cell lines) under both hypoxic and normoxic conditions were collected and lysed using RIPA buffer (Biomiga) supplemented with a protease inhibitor phenylmethanesulfonyl fluoride (PMSF) and cocktail. Proteins were separated by 10% SDS-PAGE gel followed by electrotransfer to PVDF membranes (Millipore). The membranes were blocked with TBST supplemented with 10% skim milk for 30 mins at room temperature and further probed with primary antibodies at 4°C overnight including Trim17 (1:500, Sigma); Mcl-1 (1:1000, CST); Bax (1:1000, CST); and active caspase 3 (1:500, CST) with β-actin (1:500, GeneTex) as the loading control. The membranes were subsequently incubated with HRP-conjugated secondary antibodies, and the blots were detected by ECL (Thermos Scientific). The quantifications of the protein expression levels were conducted by the gray scanning of the target proteins with Image-Pro Plus 6.0 software. Three independent experiments were performed to test the expression differences.

### shRNA-lentiviral infection and apoptosis detection of N2a cells

The HIV-derived lentiviral vector pLKO.1 containing shRNAs (TRIM17 MISSION shRNA Bacterial Glycerol Stock, Sigma) that targeted Trim17 (shTrim17-1: CCATCTGCCTTGACTACTTTA; shTrim17-2: CTGTTACCCAATTCCACTCTA) and control shRNA (shCtrl: GACACTGGGTGTGCCACAGTT) together with lentiviral packaging plasmids pCMVΔ8.9 and pMD2.G were co-transfected into HEK293T cells to generate lentiviruses. Supernatants containing different lentiviruses were collected 48- and 72-hours post-transfection, respectively. The N2a cells were infected with supernatants in the presence of 6 μg/ml polybrene for 48 hours, and 2 μg/ml puromycin was used to select the stable cell lines. Three stable cell lines expressing shRNAs were cultured simultaneously under 1% hypoxia and 21% normoxia for 48 hours and collected for FITC-conjugated annexin V (BD Pharmingen) staining followed by apoptotic cell number analysis by flow cytometry.

### Statistical analysis

Two-tailed Fisher’s exact tests were used to test the proportional differences in pseudogenes for each functional group. Paired t-tests were used to test differences in Trim17 protein levels and apoptosis rates. Hypergeometric tests (http://systems.crump.ucla.edu/hypergeometric/index.php) were used to test the significance of the overlapping pseudogenes.

## Supplementary Materials

Supplementary Table S1. Gene losses identified in each species.

Supplementary Table S2. Results of functional group enrichment analysis.

Supplementary Table S3. Mutations of pseudogenes discussed in the main text.

Supplementary Table S4. Independent gene losses between NMRs and BMRs.

Supplementary Table S5. MGI MP information for overlapping pseudogenes.

Supplementary Table S6. Vision related pseudogenes.

Supplementary Table S7. Genetic associations of pseudogenes with cardiovascular traits.

Supplementary Table S8. Functional group enrichment analysis in PSGs in NMRs and BMRs.

**Supplementary Figure S1.**
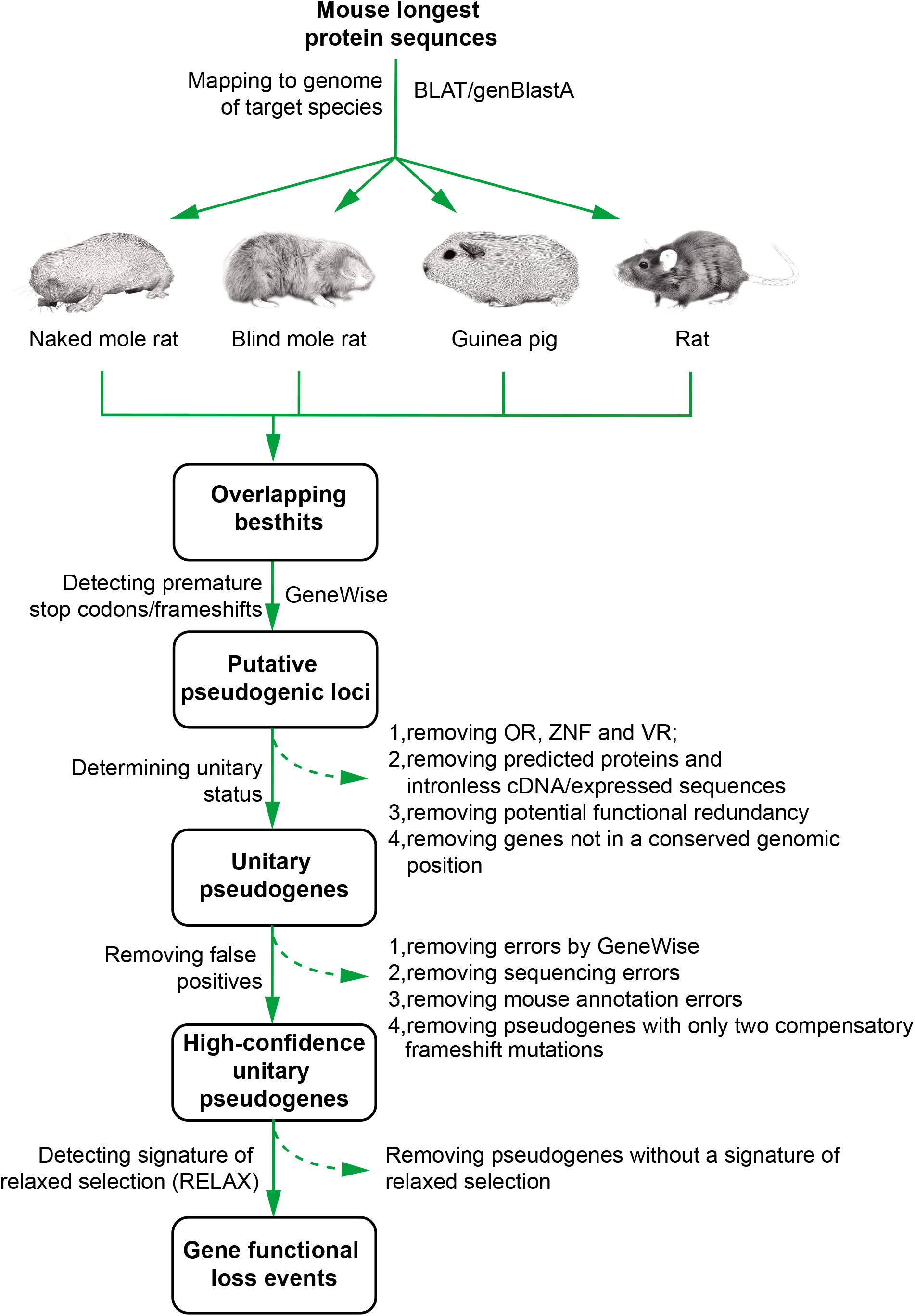
Flowchart for identifying gene loss events in NMRs, BMRs, guinea pigs and rats.

## Acknowledgments

We thank Prof. Xu-dong Zhao, Prof. Cui-ping Yang, Dr. Dong Yang and Dr. Tao Zhang for their technical support, Prof. Hua-bin Zhao for providing BMR tissue samples. We also thank Dr. Michael Hiller and two anonymous reviewers for their valuable comments. This work was supported by the Strategic Priority Research Program of the Chinese Academy of Sciences (grant no. XDB13020400), Yunnan Province, the High-End Scientific and Technological Talents program (2013HA020), and the National Natural Science Foundation of China (31871277, 31321002, 31325013, and 31301013).

